# Water-conducting roots responsible for nitrogen uptake in maize (*Zea mays*)

**DOI:** 10.1101/2024.12.28.630592

**Authors:** Yutong Jiang, Joann K. Whalen

## Abstract

Nitrate (NO_3_^-^) uptake is primarily driven by mass flow and varies among maize root types. The importance of embryonic and crown roots in acquiring NO_3_^-^ was determined here in wet and dry soils. Maize was grown in a split-root pot that segregated the embryonic and crown roots. The soil was moistened to water potentials of either -5kPa or -30kPa. A partial N mass balance was made by destructively sampling shoots, roots, and soils after 0, 24, and 48 h following ^15^N-KNO_3_ injection at the V3 and V6 stages. Gross nitrification was assessed using a ^15^N isotope dilution technique. At the V3 stage, crown roots had 202% more N uptake than embryonic roots in wet soil (-5 kPa). However, in dry soil (-30 kPa), N uptake was similar for embryonic and crown roots, possibly due to an 80% reduction of hydraulic conductance in crown roots. By the V6 stage, crown roots dominated N uptake, with embryonic roots supplying < 20% of N uptake. Soil gross nitrification rate was similar for root types. The present studies indicated that maize NO_3_^-^ uptake depends primarily on the crown roots, due to their capacity to extract water and NO_3_^-^ from soil, even under dry conditions.

**Highlight:** *This study reveals how different maize root types contribute to nitrate uptake under varying soil moisture conditions, suggesting that soil water management is important to ensure optimal nitrogen uptake*.

## Introduction

Maize (*Zea mays*) requires nitrogen (N) to produce chlorophyll and protein for maximum growth and optimal yield potential. Maize acquires more than 90% of its N supply through mass flow, primarily in the form of nitrate (NO_3_^-^) (Barber 1962; McMurtrie and Näsholm 2018). Transpiration is responsible for NO_3_^-^ movement from the soil water into the roots and then into the xylem vessels that connect the root system to the leaf surface. Continuous water flow creates tension in the xylem, producing a water gradient that draws water and associated solutes, including mobile NO_3_^-^ ions. However, the root hydraulic conductance cannot transfer NO_3_^-^ effectively from dry soil that contains decreased freely flowing water in its soil pores and breaks the continuity of water flow between the soil pores and the leaves. This issue is particularly concerning as agricultural soil water deficit is predicted to intensify under future climate conditions (Plett et al. 2020). Therefore, investigating how maize acquires NO_3_^-^ under such conditions can provide insights for soil water management, ensuring sustainable N uptake in the face of climate change.

Maize forms the primary and seminal roots during embryogenesis, and the crown and brace roots during post-embryonic development. Each root type has a unique capacity to absorb NO_3_^-^. Shoot-borne crown roots have about 63% more meta- and protoxylem elements and about 200% larger xylem vessels than embryonic primary and seminal roots (Tai et al. 2016; Hazman and Kabil 2022). This anatomical structure provides a 5-fold higher hydraulic conductance in crown roots, inducing a faster water flow from soil pores to the root xylem (Doussan 1998; Ahmed et al. 2018b). For example, the water uptake rate was 1.8 to 2.4 ×10^-5^ cm s^-1^ in crown roots and < 5 ×10^-8^ cm s^-1^ in seminal roots of a 35-d-old maize (V3 – V4 growth stage) (Ahmed et al. 2018b). An increase in hydraulic conductance can increase NO_3_^-^ uptake by the crown roots. In a hydroponic environment, the maximal NO_3_^-^ influx rate can be 22% higher in crown roots than seminal roots of 20-d-old maize (V2 – V3 stage) (York et al. 2016). This suggests that the crown roots are responsible for NO_3_^-^ uptake in the early vegetative growth (∼V3 growth stage) of maize, but it should be confirmed for a soil environment. Furthermore, if N uptake varies among the root types, then soil N transformations may vary around embryonic and crown roots due to potential differences in soil pH and the associated microbial community within adjacent soil. To the best of our knowledge, there are no published studies assessing the spatial variability of gross nitrification surrounding the maize root system.

Dry soil conditions are characterized by decreased flow of water and solutes (including NO_3_^-^) to root surfaces. The hydraulic conductance of crown roots can be decreased by 50% as the soil dries but does not change in primary and seminal roots after four weeks of water deficit (Hazman and Kabil, 2022). Recent research has revealed that the hydraulic conductivities of the seminal and primary roots decreased immediately when exposed to a mild water deficit and were followed by a full recovery in seminal roots and a 60% recovery in primary roots after 4 d of prolonged water deficit in hydroponic conditions (Protto et al. 2024). The area of xylem vessels in crown roots decreased up to 80%, while there was no change in the xylem area of primary roots (Jafarikouhini and Sinclair, 2023; Hazman and Kabil, 2022). In addition, the root may shrink and form air gaps under water deficit, which limits water flow between root-soil interfaces (Carminati et al. 2013). Crown roots may suffer from more shrinkage than seminal roots due to their 10 %–40 % thicker diameter and about 30% additional cortex cell layer (Tai et al. 2016). Consequently, embryonic roots may acquire more NO_3_^-^ than crown roots during the early vegetative growth stage under water deficit.

The objective of the present study is to quantify and compare NO_3_^-^ uptake rates by embryonic and crown roots at V3 and V6 growth stages under two contrasting soil water conditions (Ψ_soil_ = -5 and -30 kPa) using a customized split-root system combined with the ^15^N isotope technique. It is hypothesized here that crown roots have a higher NO_3_^-^ uptake rate than embryonic roots in relatively wet soil (soil water potential Ψ_soil_ = -5 kPa) at both V3 and V6 growth stages due to their higher hydraulic conductance. However, embryonic roots in relatively dry soil (Ψ_soil_ = -30 kPa) are expected to acquire more NO_3_^-^ than crown roots at the V3 growth stage because crown roots have decreased hydraulic conductance when exposed to the dry soil. A higher nitrification around crown roots than embryonic roots in the relatively wet soil is also anticipated.

## Materials and Methods

### Soil and maize

Soil (0 –15 cm) was collected from the Emile A. Lods Agronomy Research Centre in Sainte-Anne-de-Bellevue, Québec (45°25′N, 73°55′W, 39 m elevation) in October 2022 following maize harvest. The soil is a sandy-loam Humic Gleysol of the St. Amable series containing 620 g sand kg^-1^, 60 g clay kg^-1^, and 48 g organic C kg^-1^, with a pH of 6.3. Each kg of soil (sieved < 6 mm) was pre-mixed with NPK fertilizers (expressed as %) containing 37 mg urea (46-0-0), 59 mg triple superphosphate (0-46-0), and 28 mg potassium chloride (0-0-60). The maize variety was *Zea mays* L. cv. PS2790, which was neither genetically modified nor treated with a fungicide or insecticide. All chemicals and reagents were purchased from commercial sources and were at least analytical grade purity.

### Split-root pot design

The experimental split-root pot was designed to segregate the embryonic root system (i.e., primary and seminal roots) from the crown roots (Fig. S1). An inner chamber (PVC pipe with 5 cm diameter, 25 cm height) was placed in the middle of the outer chamber (PVC pipe with 10 cm diameter, 30 cm height). Field-moist soil was packed into the inner chamber (0.54 kg dry weight basis) and the outer chamber (2.05 kg dry weight basis), with a constant bulk density of 1100 kg m^-3^. An aluminum foil cone (3 cm height) was placed on top of the inner chamber. A small hole (0.5 cm dia.) was pierced on the top of the cone, allowing the primary and seminal roots (i.e., the embryonic roots) to grow through the hole into the inner chamber, while the crown roots are directed to grow in the outer chamber. There was no drainage in the split-root pot to avoid water and soluble nutrient losses by leaching.

### Experimental treatments

A two-way factorial experimental design was used to evaluate the N acquisition by maize as influenced by soil moisture (-5 kPa or -30 kPa, measured as soil water potential, Ψ_soil_) and the maize root type (embryonic roots or crown root), with four replicates per treatment. The maize root types were separated spatially within the split-root pot, and each pot was a replicate in the experiment. The soil water potential was monitored in eight additional split-root pots at two water levels (n = 4 for each water level) with a Model MLT tensiometer (Irrometer Inc., Riverside, California, US) inserted into the inner chamber and the outer chamber (Fig. S2). Additional control studies were conducted to confirm that the maize root growth in a split-root pot was similar to a regular PCV pot (10 cm diameter, 30 cm height) by growing maize in eight pots (= 2 water levels × 4 replicates) each containing 2.4 kg soil (dry weight basis) and no inner chamber (Fig. S3).

Maize seeds were pre-germinated in pots (12.7 cm dia., 15 cm depth) for 10 d before transplanting to the split-root pots. Embryonic roots emerged within 10 d after seeding, but crown roots did not. Thus, embryonic roots were manually directed into the inner chamber through the small hole on the aluminum foil cone, while the crown roots emerged 3 to 5 d later and grew naturally into the outer chamber. All pots were placed in a growth bench under a fluorescent lighting system (400 µmol^-2^ s^-1^ of photon flux density) with controlled light and temperature of 16 h light (25 ± 2 °C) and 8 h dark (18 ± 1 °C). Distilled water was added every 2 d using a syringe with a 12-cm long needle (HaBeuniver Inc., Fuzhou, Fujian, China) to maintain the soil water potential at target levels (Ψ_soil_ = -5 or -30 kPa) according to the average tensiometer readings in the eight monitoring pots.

At the V3 and V6 growth stages, N acquisition by each maize root type was evaluated using ^15^N-labeled KNO_3_. When maize had three fully-formed leaves (V3), KNO_3_ solution (0.33 g N L^-1^, 18.4% atom ^15^N excess) was injected either in the inner (26 mL) or the outer chamber (112 mL) of the pots, targeting at Ψ_soil_ = -5 kPa. A different KNO_3_ solution (0.67 g N L^-1^, 18.7% atom ^15^N excess) was injected either in the inner (13 mL) or the outer chamber (56 mL) of the pots that targeted at Ψ_soil_ = -30 kPa to deliver 0.02 g N kg^-1^ soil. A non-labeled KNO_3_ solution (0.33 g N L^-1^ or 0.67 g N L^-1^) was injected into the chamber that was not treated with the ^15^N-KNO_3_ solution, so the total N input was the same for the inner and outer chambers of each pot. Controlled pots were handled identically except that both inner and outer chambers received non-labeled KNO_3_ solution to establish the ^15^N natural abundance in maize. Pots (n = 56) were destructively sampled at 0, 24, and 48 h post-labeling. The pots that were kept until the V6 stage (n = 56) received ^14^N-KNO_3_ in the inner and outer chambers at the V3 stage. At the V6 growth stage, these pots were labeled and sampled following the procedure described above for the V3 growth stage.

### Plant sampling and analysis

Following exposure to ^15^N-KNO_3_, pots were destructively sampled by cutting maize stems at the soil surface, then the shoots were rinsed, dried (55 °C for 72 h), and ground (< 1 mm). Roots were removed from the soil by gently shaking and removing most of the root-associated soil. A single crown root, seminal root, and primary root from each pot were immediately removed under deionized water for root hydraulic conductance analysis. The rest of the roots were washed thoroughly with deionized water to remove the attached soil particles and then rinsed with 1 mM CaSO_4_ to remove apoplastic ^15^N. Samples were stored in a refrigerator at 4 °C for up to 7 d, while the following analyses were underway.

Root hydraulic conductance was measured within 1 h of sampling by the root exudation method (Knipfer et al. 2010). The single primary root, seminal root, and crown root were connected to a glass microcapillary tube (0.3-, 0.5-, or 1-mm diameter). Then, the roots sampled from pots with Ψ_soil_ = -5 or -30 kPa were bathed in a 0.069 or 4 g L^-1^ sucrose solution. The movement of exudate in the capillary was recorded every 15 min for 3 h and the water flow rate (mm^3^ min^-1^) was calculated as the slope of the linear part of the flow vs time plot. After 3 h, the exudate was collected from the capillary tube using a syringe needle. The osmotic pressures of the bath medium and the xylem exudates were determined by measuring the osmolality using a Model 5520 vapor pressure osmometer (Wescor Vapro, Utah, USA). The surface area (m^2^) and length (m) of the single root used for the hydraulic conductance measurement was determined using a Model V700 flatbed photo scanner (Epson Inc., Markham, Canada) and WinRhizo software (Regent Instruments, Quebec, Canada). The root hydraulic conductance (m^3^ s^-1^ MPa^-1^) was calculated using Equation (1):

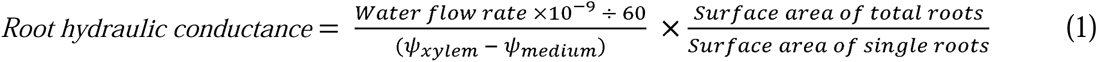

where, ψ*_medium_* is the osmotic pressures of the bath medium (MPa), ψ*_xylem_* is the osmotic pressures of the xylem exudates (MPa).

Hand cross-sections of the tip (0 – 20 mm from root tip), mid (50 – 70 mm from root tip), and base (0 – 20 mm from root base) of the crown root, seminal root, and primary root were prepared for anatomical analyses. Root segments were fixed in 4% paraformaldehyde in 1× phosphate buffer saline solution (PBS) while shaking (3 h, 22 °C), and washed 3 times (30 min, 22 °C) in 1× PBS. The root segments were thin-sectioned, stained with 0.1% Toluidine Blue O, and washed with 1× PBS solution before sealing with a cover slip. Root anatomy images were taken with light microscopy (ZEISS imager Z1, Germany) with ×10 magnification. The xylem area, number, diameter, stele area, and cross-section area were analyzed by ImageJ (Fiji version for Mac OS X, NIH, Bethesda, MD, USA). The cortex area was calculated by subtracting the stele area from the cross-section area. The hydraulic conductance of tip (0 – 20 mm from root tip), mid (50 – 70 mm from root tip), and base (0 – 20 mm from root base) segments were calculated (Equation 2) using the Hagen-Poiseuille Law (Strock et al. 2021) expressed as

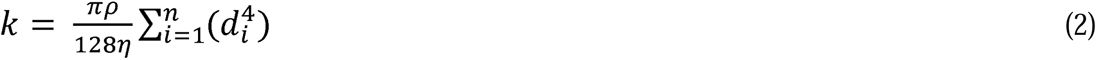

where ρ is the xylem sap fluid density assumed to be water (1000 kg m^-3^, at 20 °C), and η is the viscosity of the xylem sap assumed to be water (1 × 10^-9^ MPa s, at 20 °C), and d is the mean diameter of each metaxylem vessel (m).

Morphological traits (length and surface area) of primary, seminal, and crown roots were measured by WinRhizo software. Total root length and total surface area also included the excised roots subsampled for root hydraulic and anatomy measurements. The length and surface area of lateral roots were calculated as the sum of the fine roots with a diameter < 0.5 mm as measured using WinRhizo software.

Shoots, embryonic roots, and crown roots were oven-dried (at 55 °C for 48 h), weighed (for dry mass), and ground (< 1 mm). The N content and ^15^N excess (%) of the shoot (*[APE]_shoot_*), primary root (*[APE]_primary_*), seminal (*[APE]_seminal_*), and crown root (*[APE]_crown_*) were measured using 2 – 5 mg of root tissue analyzed by an isotope ratio mass spectrometer at the UC Davis Stable Isotope Facility (Davis, CA, USA). The *N* uptake rate (mg d^-1^) was calculated (Equation 3) according to Barraclough (1996):

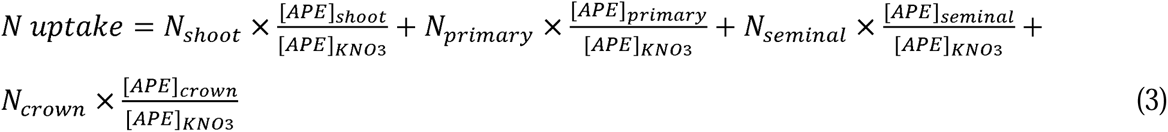

where, *[APE]_shoot_*, *[APE]_primary,_ [APE]_seminal,_* and *[APE]_crown_* are N content and ^15^N excess (%) of the shoot, primary roots, seminal roots, and crown roots. N_shoot_, N_primary_, N_seminal_, N_crown_are N content of shoot, primary roots, seminal roots, and crown roots.

### Soil sampling and analysis

Soil from the inner chamber and the outer chamber were removed separately and stored at 4 °C for up to 7 d until analysis. Soil NH_4_ ^+^ and NO_3_^-^ concentrations were analyzed in extracts from 20 g soil mixed with 80 mL 0.5M K_2_SO_4_ solution (1:4, soil: 0.5M K_2_SO_4_ solution) and measured calorimetrically at 650 nm on a µQuant microplate reader (BioTek, Winooski, VT, USA) following Sims et al. (1995),

The ^15^N enrichment of NH_4_ ^+^ and NO^3-^ pools was determined using an acid diffusion procedure (Brooks et al., 1989), with > 90% recovery of NH_4_ ^+^ and NO_3_^-^ from the extracts based on internal standard analysis. Acidified (15 µL of 2.5 M KHSO_4_) glass filter paper disks (5 mm, Whatman GF/D, pre-ashed at 500 °C for 4 h) were sealed in Teflon tape (2 disks per Teflon packet). To attain the required 20 – 50 µg N in solution and considering the low NH ^+^ concentration (< 2 mg NH_4_ ^+^ kg^-1^) in soil extracts, ten mL of soil extract was spiked with 10 mL ^15^N-(NH_4_)_2_SO_4_ (4 mg N L^-1^, 4.2 atom % excess ^15^N) in a 120-mL specimen cup before the diffusion procedure. The ^15^N enrichment of NO_3_^-^ pool was evaluated after diluting the soil extract with 10 mL of 0.5M K_2_SO_4_ in a separate 120-mL specimen cup. Diffusion began after adding one Teflon packet and 0.1 g MgO (for ^15^N-NH_4_) or one Teflon packet, 0.1 g MgO, and 0.4 g Devarda’s alloy (for ^15^N-NO_3_) and capping the specimen cup. Blanks (K_2_SO_4_) and the spiking solution (^15^N-(NH_4_)_2_SO_4_) were also analyzed to correct for the background ^15^N concentration and ^15^N recovery, respectively. Specimen cups were mixed gently by hand twice a day for 7 d to ensure the reaction of MgO and Devarda’s alloy with the sample solution. After 7 d, filter paper disks were removed from the Teflon packet, dried over concentrated H_2_SO_4_ in a desiccator for about 48 h, and packaged in tin capsules for isotopic enrichment or atom percent enrichment (APE) determination on a Micro Cube elemental analyzer (Elementar Analysensysteme GmbH, Hanau, Germany) interfaced to a PDZ Europa 20-20 isotope ratio mass spectrometer (Sercon Ltd., Cheshire, UK) at the UC Davis Stable Isotope Facility.

### Nitrogen calculations

The atom % ^15^N in the NH_4_-Npool was calculated using Equation (4):

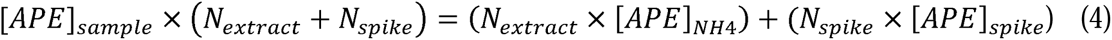

where, *[APE]_sample_* is the atom % ^15^N measured from the diffusion disks, *N_extract_*is the amount of NH_4_-N (µg) in soil extracts, *[APE]_spike_* is the atom % ^15^N measured from the diffusion disks from the spike solution, and *N_spike_* is the amount of NH_4_-N (µg) in the spike (Whalen et al. 2001). The unknown in the equation is *[APE]_NH4_*, the atom % ^15^N in the NH_4_-N pool of soil extracts.

Similarly, atom % ^15^N in the NO_3_-N pool was calculated using Equation (5):

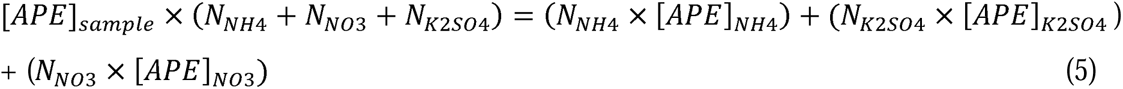

where *N_NO3_* is the amount of NO_3_-N (µg) in soil extracts, *N_K2SO4_* is the amount of N in the blank solution, and *[APE]_K2SO4_* is the atom % ^15^N measured from the diffusion disks from the blank solution. The unknown in the equation is *[APE]_NO3_*, the atom % ^15^N in the NO_3_-N pool of soil extracts.

Gross nitrification rate (mg of N kg^-1^ soil d^-1^) was calculated using Equation 6 (Kirkham & Bartholomew, 1954):

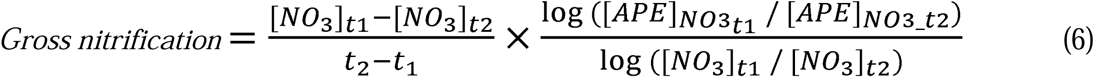

Where *[NO_3_]_t1_* and *[NO_3_]_t2_*represent the total NO_3_^-^ concentration (mg N kg^-1^) at *t_1_* (time *1,* 0 d) and *t_2_* (time *2,* 7 d), respectively. *[APE]_NO3_t1_* and *[APE]_NO3_t2_* indicate atom % ^15^N excess of NO_3_^-^ at *t* and *t,* respectively.

### Statistical analyses

Data was normally distributed (Shapiro-Wilk test, *P* > 0.05) with homogeneous variance (Levene’s test, *P* > 0.05). Analysis of variance revealed no effect (*p* >0.05) of sampling time (0, 24, and 48 h post-labeling) on several dependent variables (root length and root hydraulic conductance). Thus, data were pooled among sampling times, which gave more replicates for root morphological measurements (n = 20, Fig. 1) and root hydraulic measurements (n = 12, Fig. 2) for each treatment. The main and interactive effects of root types and soil water potential were evaluated by analysis of variance. When the main effects or interactive effects were significant (*p* < 0.05), mean values were compared with Tukey’s honestly significant test. The xylem number, xylem area, and calculated hydraulic conductance under the two water potentials were compared by Student’s *t*-test. Correlation with the Pearson coefficient was used to describe the best-fit lines of the relationship between root hydraulic conductance of each root type and N uptake (R 3.6.1).

**Fig. 1.**
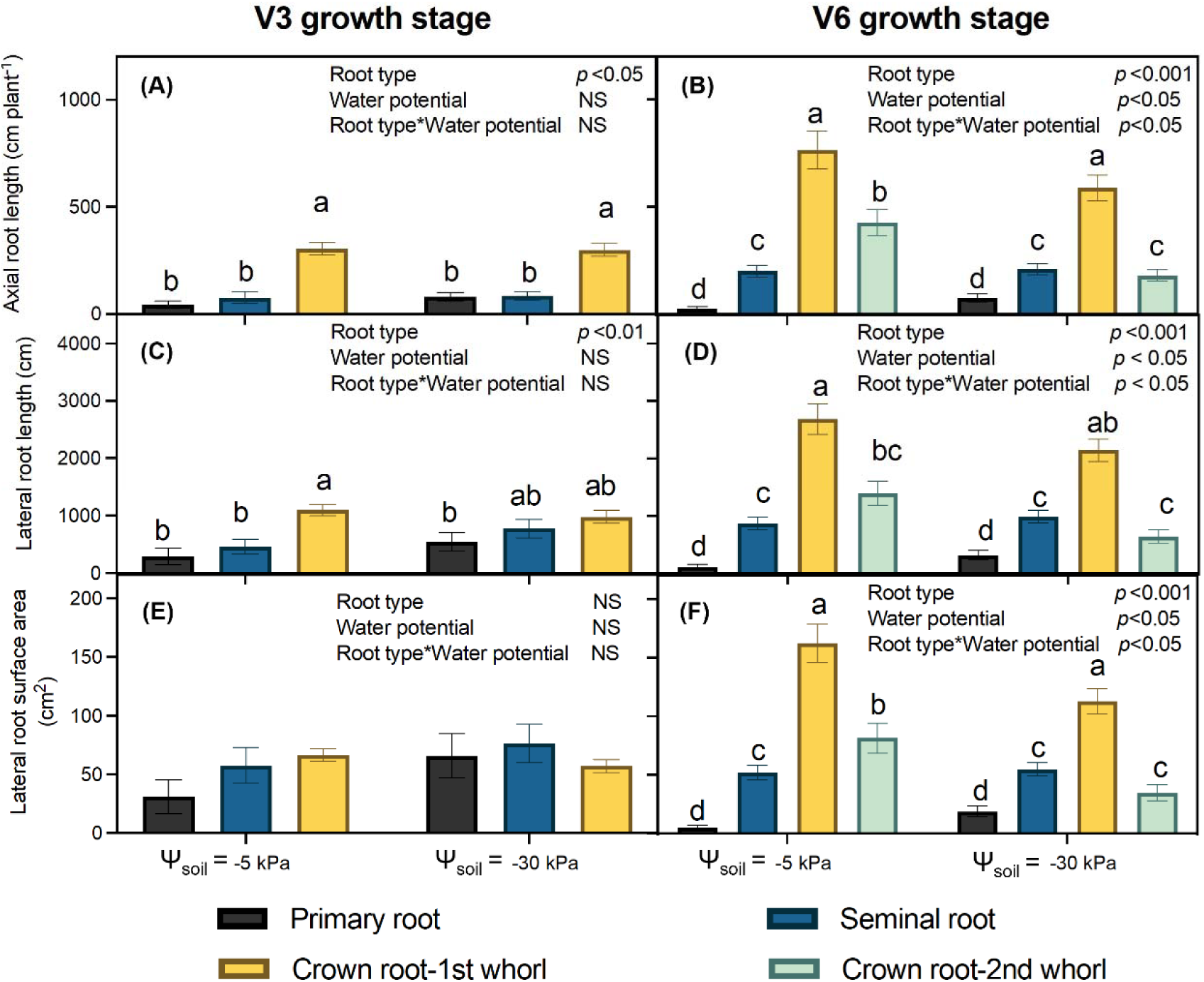
Root morphological traits. Panels (A, B) axial root length; (C, D) lateral root length, and (E, F) lateral root surface area of maize grown in the split-root system to the V3 or V6 stages. Maize was grown in pots with a constant water potential (-5 kPa or -30 kPa). Data was pooled among N tracer treatments, and plants with the same water potential were pooled together (n = 20). Different letters over the bars represent significant differences (*p* < 0.05, Tukey’s honestly significant test). NS, not significant (*p* > 0.05).

**Fig. 2.**
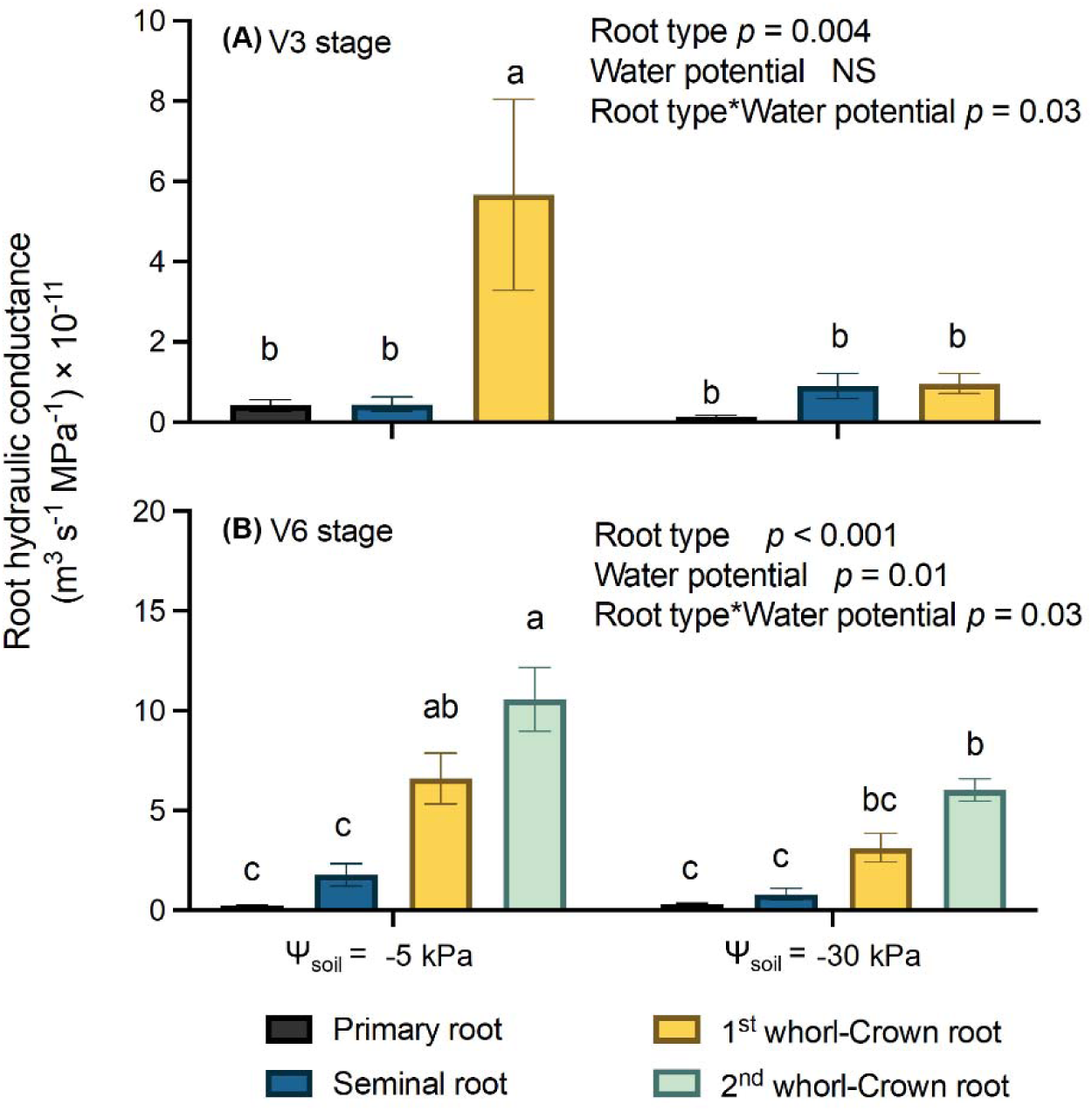
Measured hydraulic conductance of maize grown in the split-root system. Panel (A) shows hydraulic conductance at V3 and Panel (B) at V6 stages. Maize was grown in pots with a constant water potential (-5 kPa or -30 kPa). Hydraulic conductance was determined with the root exudation method, with one primary, seminal, 1^st^ whorl crown root, and 2^nd^ whorl crown root collected from each of the 12 plants grown under one water potential for analysis (n = 12). Error bars represent the standard error of means. Different letters over the bars represent significant differences (*p* < 0.05, Tukey’s honestly significant test). NS, not significant (*p* > 0.05).

## Results

### Morphology of primary, seminal, and crown roots

By the V3 growth stage, crown roots emerged as the predominant root type, constituting 65– 72% of total axial roots (Fig. 1). The lateral root length was 2 – 3 times greater on the crown than on the primary and seminal roots when exposed to the wet soil (Ψ_soil_ = -5kPa) (Fig. 1c). However, in dry soil (Ψ_soil_ = -30kPa), all root types produced lateral roots having similar lengths and surface areas of (*p* > 0.05; Fig. 1c, e). At the V6 stage, the crown root system, comprising of 1^st^ and 2^nd^ whorl crown roots, constituted 73–84% of axial root length and 68–81% of lateral root length. Lateral surface area was 300 – 328% greater for crown roots than embryonic roots, regardless of soil water potential. When exposed to dry soil, the 2^nd^ whorl crown roots produced 58% shorter axial roots and 57% less lateral root surface than grown in the wet soil.

### Effect of dry soil on hydraulic conductance of crown roots

At the V3 growth stage, crown roots had 22% more xylem vessels and 56 – 120% larger xylem area than primary roots, and 75% more xylem vessels and 84 – 300% larger xylem area than seminal roots in wet soil (Table 1). The increased xylem vessel area was associated with the 80 – 600 % higher calculated hydraulic conductance along the crown roots than along the primary and seminal roots. Dry soil had 1^st^ whorl crown roots with 35% less xylem area at the root tip, leading to 46% lower hydraulic conductance in this region. In contrast, the hydraulic conductance was similar in primary and seminal roots growing in the dry soil (*p* > 0.05). At the V6 stage, primary and seminal roots contributed < 5% of hydraulic conductance, regardless of soil water condition. The 1^st^ and 2^nd^ whorl crown roots had 16 – 45 times higher calculated hydraulic conductance than the primary and seminal roots. The 2^nd^ whorl crown roots had 13% reduced xylem area and 25% reduced root hydraulic conductance at the root tip region. Measured root hydraulic conductance revealed a consistent pattern among different root types (Fig. 2). At the V3 stage, crown roots had 10 times more hydraulic conductance than primary and seminal roots in wet soil (*p* < 0.05).

**Table 1.**
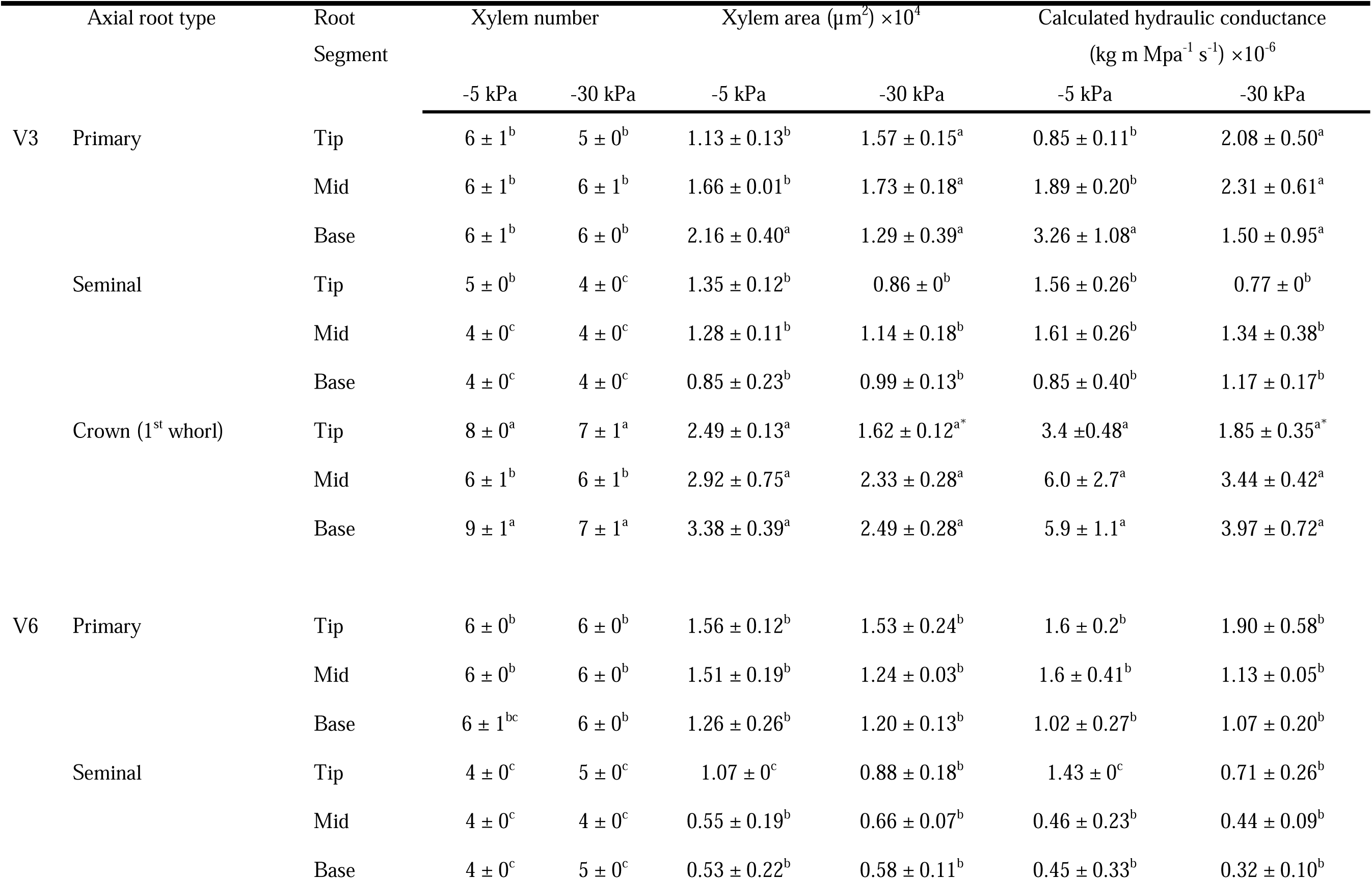
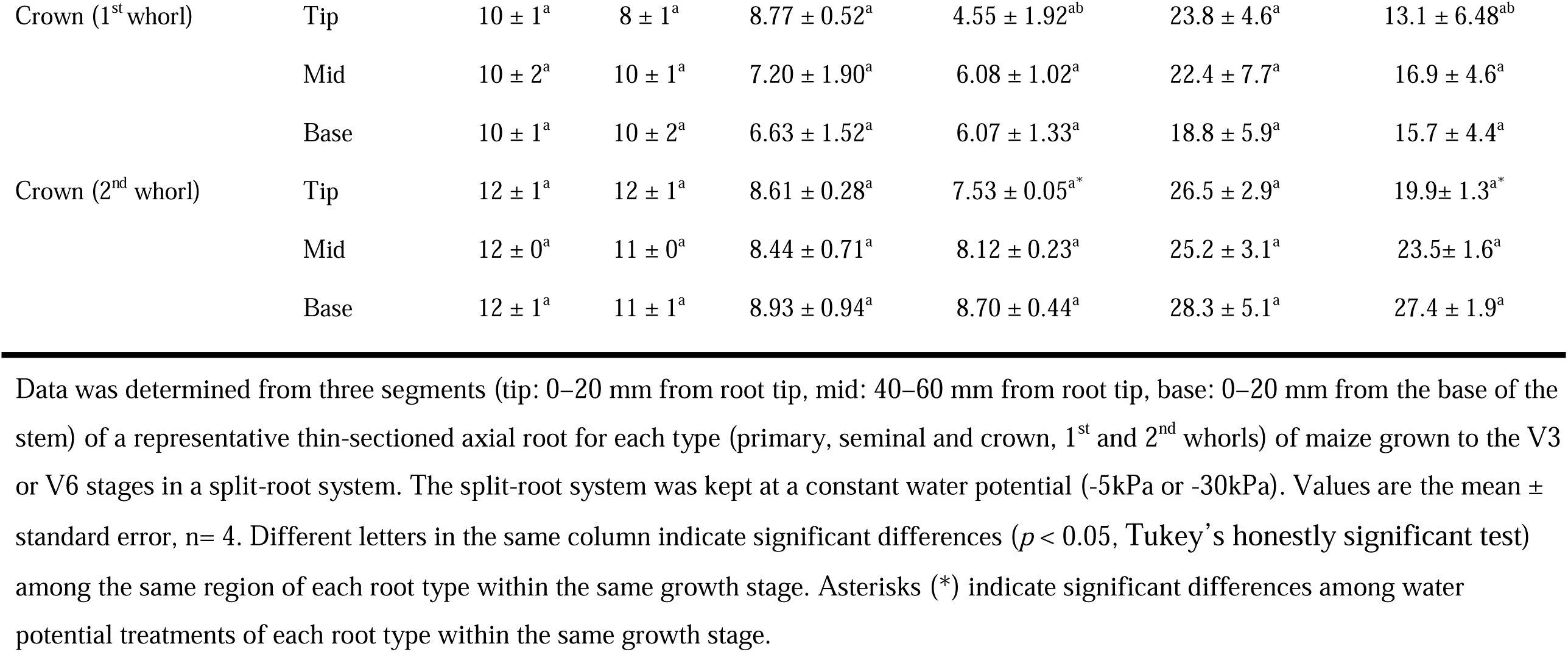
Xylem number, xylem area, and calculated hydraulic conductance.

However, the hydraulic conductance of crown roots decreased by 83% in dry soil because root hydraulic conductance interacts with soil water potential (Root type × Water potential effects, *p* = 0.03). Similar hydraulic conductivities in crown, primary, and seminal roots in the dry soil were observed. By the V6 stage, the 1^st^ and 2^nd^ whorl crown roots dominated and contributed to 90% of water-conducting capacity, despite a 37% reduction in 2^nd^ whorl crown root hydraulic conductance in dry soil.

### Effects of dry soil on N uptake by the crown and embryonic roots

Between 80% to 112% of the ^15^N was recovered in the soil, shoot, and root pools from 0, 24, and 48 h after injection of the ^15^N-KNO_3_ solution at V3. Similarly, between 83% to 118% of the ^15^N was recovered in the soil, shoot, and root pools up to 48 h after injection of the labeled solution at V6 (Table 2). Maize shoot and root biomass were progressively enriched with ^15^N in the hours after injection of the ^15^N tracer (Fig. S4). At the V3 growth stage, N uptake by crown roots was 2.7-fold higher than embryonic roots in the wet soil (*p* < 0.05, Fig.3). However, in the dry soil, the N uptake rate was similar in crown and embryotic roots (*p* > 0.05), with embryonic roots supplying 38% of N uptake. By the V6 stage, the crown root system was the dominant pathway for N uptake that was responsible for about 80% of N uptake by maize growing in wet and dry soils.

**Fig. 3.**
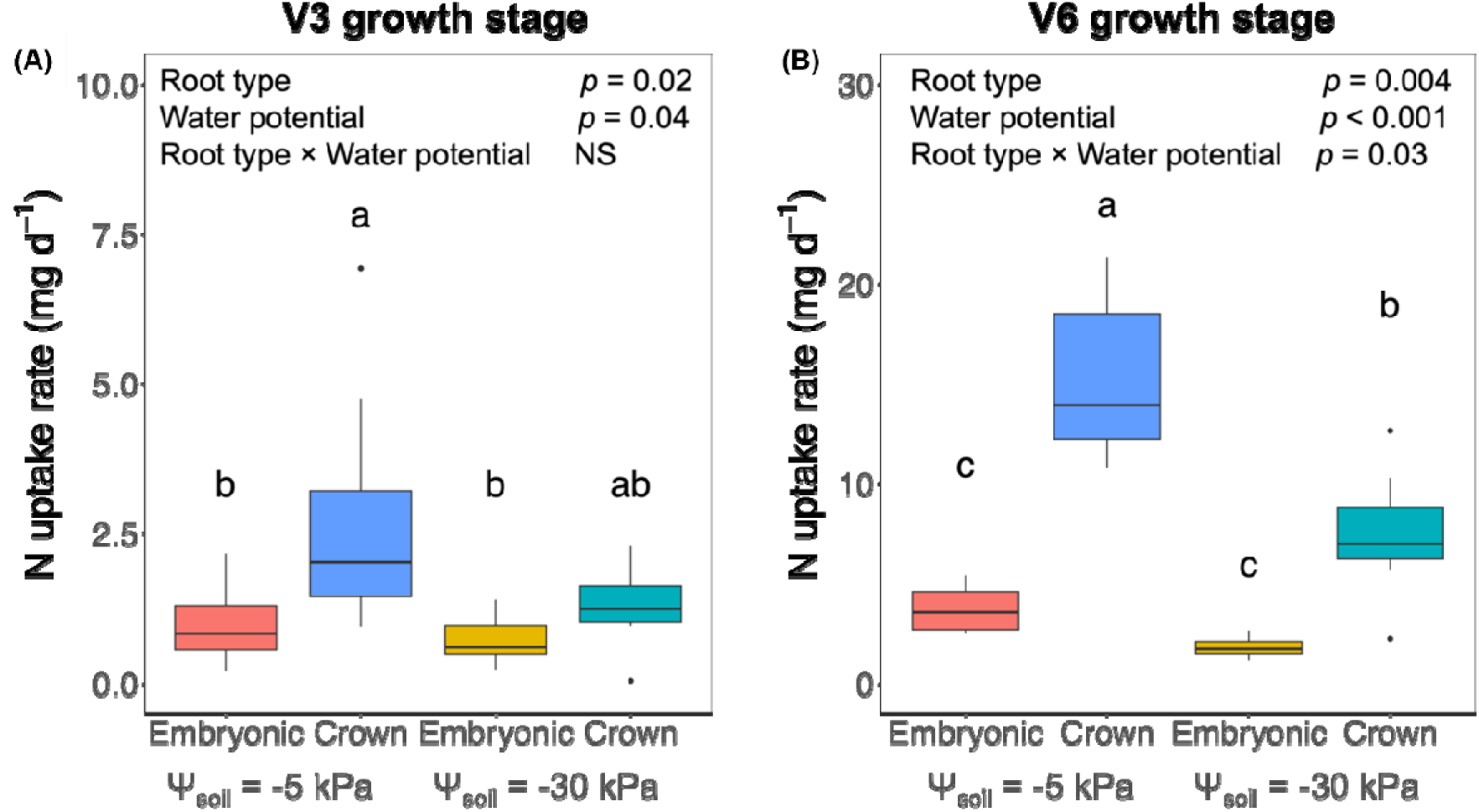
Nitrogen uptake by embryonic roots and crown roots of maize grown in the split-root system. Panel (A) shows N uptake during the V3 stage, and Panel (B) shows N uptake during the V6 stage. Maize was grown in pots with a constant water potential (-5 kPa or -30 kPa). Nitrogen uptake rate was calculated from the ^15^N enrichment in maize shoot after 0, 24, and 48 h ^15^N enrichment periods. At the V3 and V6 stages, one of the chambers, either the inner or outer, received an injection of ^15^N-KNO_3_ solution (18.4 % atom ^15^N excess), while the other chamber received ^14^N-KNO_3_ solution, ensuring exposure of only one root type (either embryonic or crown roots) to ^15^N-KNO_3_ solution. Boxplots show minimum, median, and maximum. The dots show outliers. Different letters over the bars indicate significantly different (*p* < 0.05, Tukey’s honestly significant test). NS, not significant (*p* > 0.05).

**Table 2.**
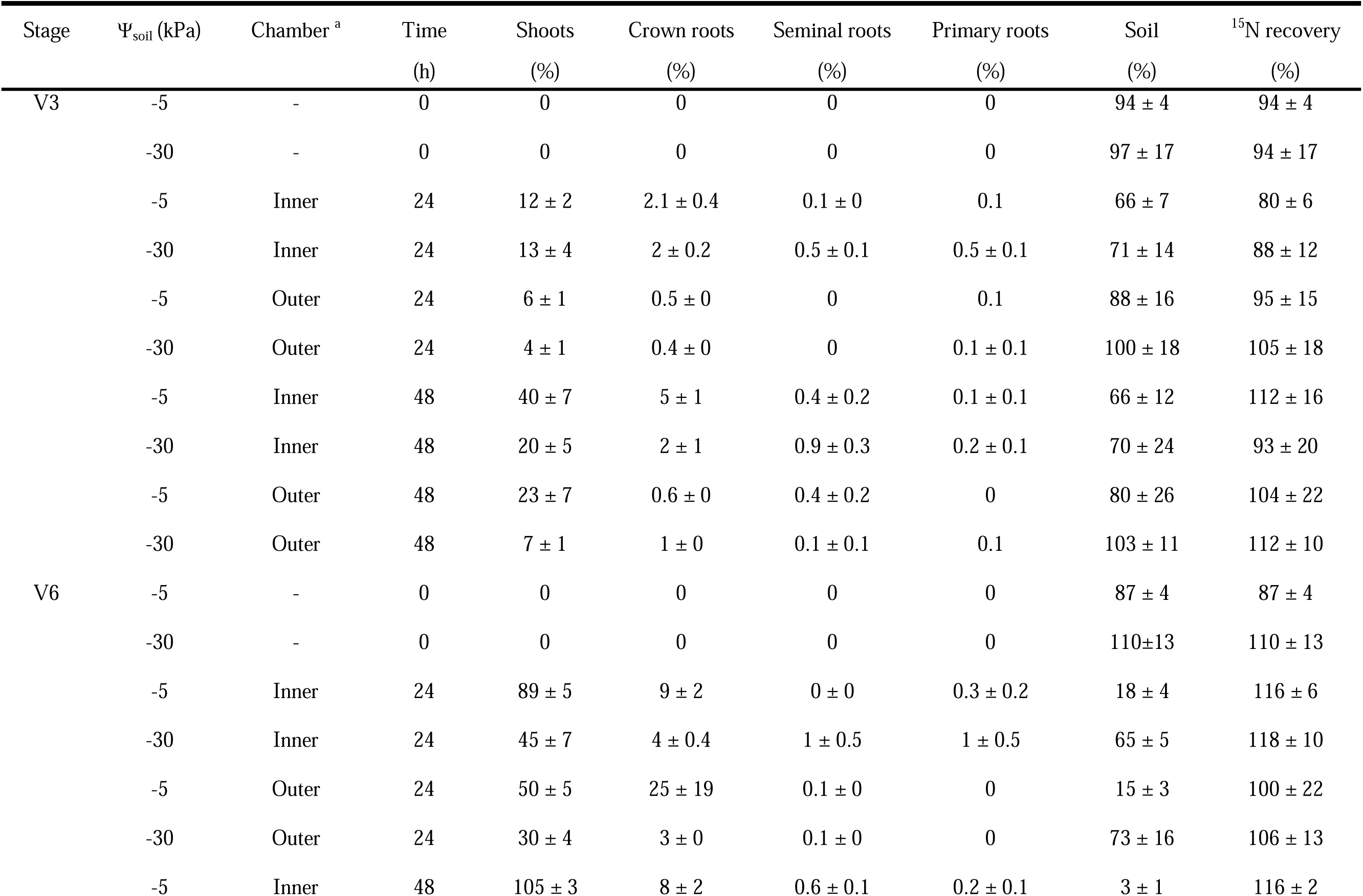
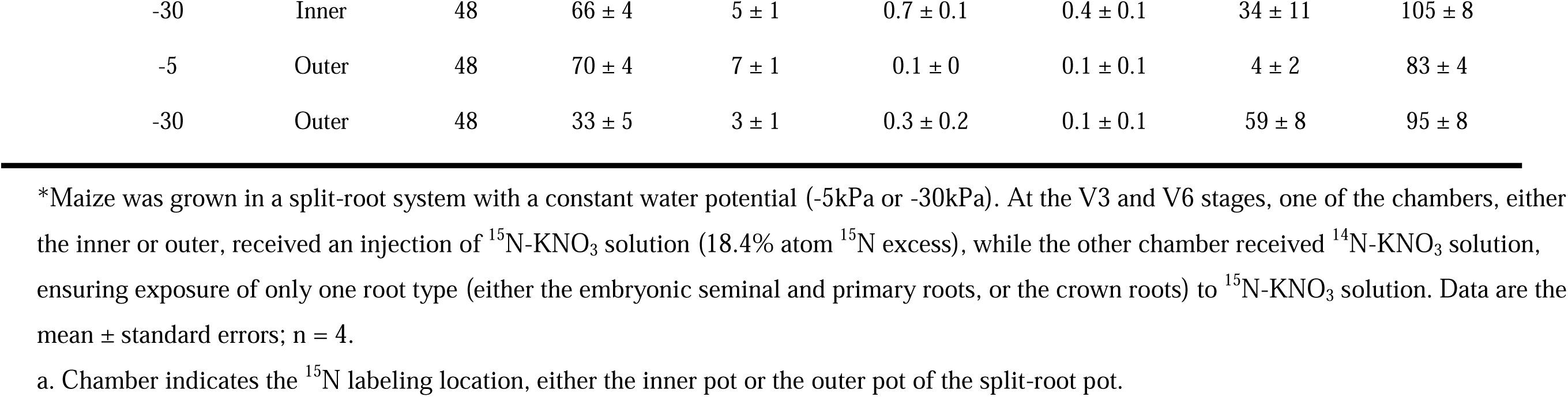
Percentage of ^15^N from added ^15^NO_3_-N in maize shoot, roots, and soil after 0, 24, and 48 h of exposure to ^15^N enrichment*.

### Nitrification among the root types

Gross nitrification rates were similar at the V3 and V6 growth stages, ranging from 3.8 – 45.5 mg NO_3_^-^ kg^-1^ d^-1^ (Fig. 4). Root types did not affect the nitrification rate at the V3 and V6 growth stages. The nitrification rate was 330% higher around crown roots in the wet soil than in the dry soil at the V6 growth stage.

**Fig. 4.**
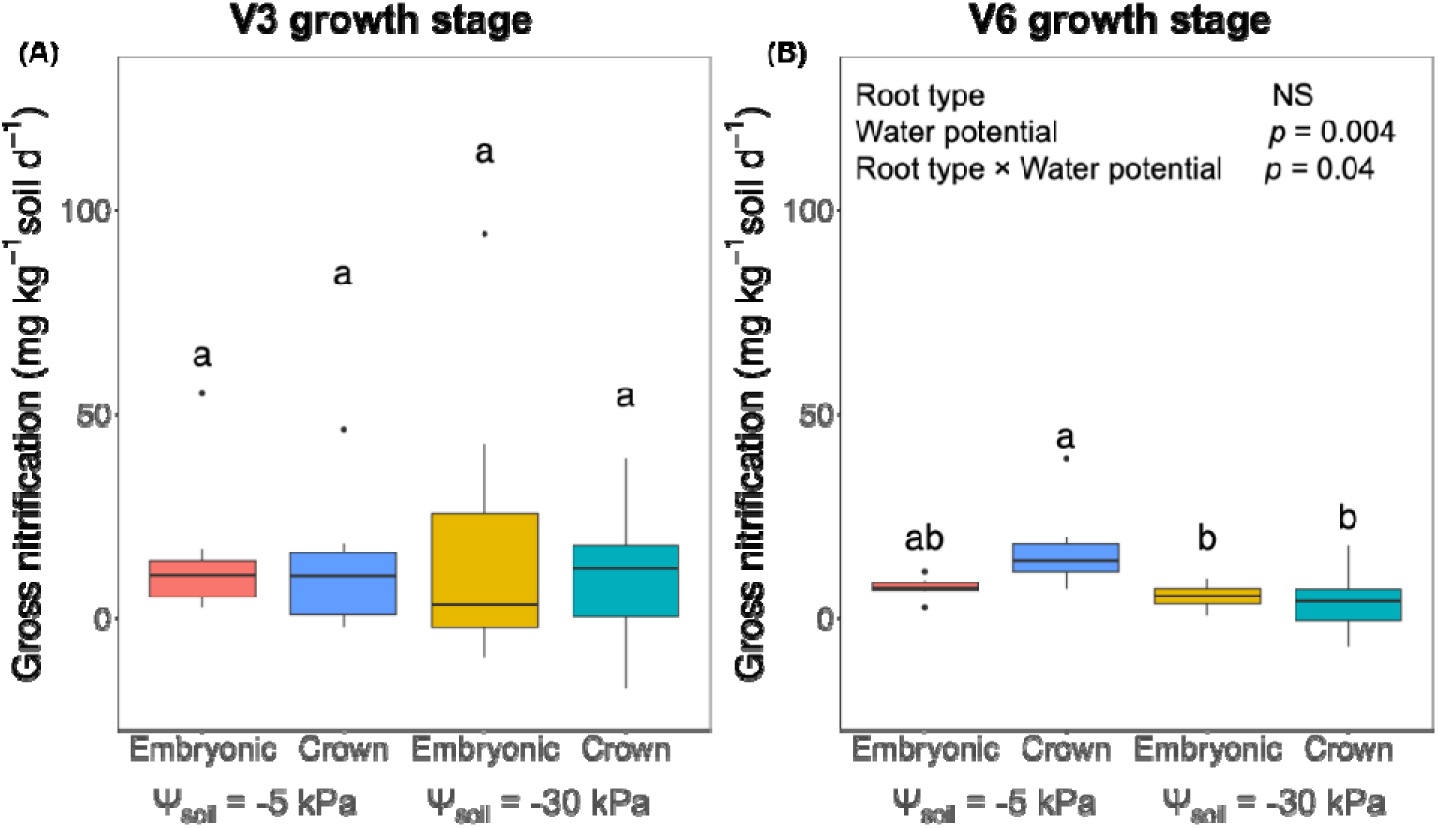
Gross nitrification around embryonic roots and crown roots of maize grown in the split-root system. Panel (A) shows gross nitrification at the V3 stage and Panel (B) at the V6 stage. Maize was grown in pots with a constant water potential (-5 kPa or -30 kPa). The gross nitrification rate was calculated from the ^15^N isotope dilution after 0, 24, and 48 h ^15^N enrichment periods. At the V3 and V6 stages, one of the chambers, either the inner or outer, received an injection of ^15^N-KNO_3_ solution (18.4% atom ^15^N excess), while the other chamber received ^14^N-KNO_3_ solution, ensuring exposure of only one root type (either embryonic or crown roots) to ^15^N-KNO_3_ solution. Boxplots show minimum, median, and maximum. The dots show outliers. Different letters over the bars indicate significantly different (*p* < 0.05, Tukey’s honestly significant test). NS, not significant (*p* > 0.05).

## Discussion

The present study tests the hypothesis that crown roots have a higher NO_3_^-^ uptake rate than embryonic roots in relatively wet soil at both V3 and V6 growth stages. Data indicated that this hypothesis could not be rejected and therefore was accepted (Fig.3). In the present study, NO_3_^-^ was the source of plant-available N for maize because NO_3_^-^ represented more than 95% of the soil mineral N pool (Fig. S5). Thus, it can be assumed that mass flow was the dominant process for maize N uptake, and NO_3_^-^ moved in the water that was extracted from soil pores by maize roots. Data indicated that NO_3_^-^ was mainly taken up by crown roots in wet soil (-5 kPa), at both V3 and V6 growth stages (Fig.3). This is consistent with other studies that the water uptake rate was 1000 times greater in crown roots than in seminal roots of 35-day-old maize (Ahmed et al. 2018b), suggesting a higher mass flow capacity in crown roots. Our observation also aligns with Liu et al. (2020), who reported that crown roots of barley (*Hordeum vulgare*) had 2-fold higher NO_3_^-^ uptake and root-to-shoot translocation capacities than the seminal roots. Despite genetic differences between barley and maize, both cereals share similarities in their root systems, characterized by a typical fibrous root system comprising embryonic seminal roots and post-embryonic crown roots. By the V6 stage, a five-fold increase in plant NO_3_^-^ uptake was observed compared to the V3 stage (Fig.3), suggesting that maize enters the rapid N uptake phase at V6 (Bender et al. 2013). The crown root system dominates the N uptake in both wet and dry soil, accounting for about 80% of N uptake by maize. This result aligns with the fact that the contribution of embryonic roots decreases as maize grows beyond the V3 stage (Ahmed et al. 2016, 2018b).

The second hypothesis that the embryonic roots had higher N uptake than crown roots in dry soil was rejected. Yet, the contribution of the embryonic root system to plant N uptake at the V3 growth stage increases from 25% to 38% as the soil dries (Fig.3). The results presented here indicate a positive correlation between NO_3_^-^ uptake and the root hydraulic conductance of shoot-derived crown roots (Fig. S6), aligning with the transport mechanism of NO_3_^-^ ions (McMurtrie and Nasholm, 2018). This relationship suggests that increased hydraulic conductance in crown roots facilitates the transport of NO_3_^-^ to roots (Henriksson et al. 2021). However, the root hydraulic conductance of crown roots was decreased by 83% when exposed to dry soil (Table 1, Fig. 2), while primary and seminal roots maintained their root hydraulic conductance. This result is consistent with Hazman and Kabil, (2022), who reported that crown root hydraulic conductance had a 50% reduction, while embryonic root hydraulic conductance was insensitive when exposed to the water-deficit. The reduced root hydraulic conductance decreased N uptake, as a 52% reduction of N uptake by crown roots in the dry soil was observed at V6 (Fig.3), which is consistent with other studies showing a reduction in N uptake by maize in response to decreased soil water availability (Hammad et al., 2017; Flynn et al., 2023). Given the significant impact of the interaction between water potential and root types on N uptake at the V6 stage, the results reported here underscore the critical importance of soil water management during this growth stage to optimize both water and N uptake in maize.

There are several explanations for the reduction of root hydraulic conductance in crown roots. This reduction was not related to the decreased root length or surface area under water deficit, as only the root morphology of 2^nd^-whorl crown roots responded to soil water potential (Fig. 1). Instead, it is likely due to the plastic responses of xylem number and diameter in the crown root tip under water deficit. The diameter of 1st whorl crown roots and 2nd whorl crown roots decreased from 64 to 58 µm and 96 to 92 µm (Table 1), which is smaller in magnitude than the reductions observed in maize (cv. CML538) root tips, whose diameter decreased from 150 to 30 µm (Jafarikouhini and Sinclair, 2023). The xylem vessel diameter and their plasticity to water deficit vary among maize genotypes, with a wide range of diameters (from 50 – 150 µm) among 44 genotypes (Yang et al. 2019), and only two out of four genotypes showed reduced metaxylem number and area under water deficit (Hazman and Kabil et al. 2022). These findings suggest that less plastic genotypes may have the potential to maintain their N uptake under water deficit, which needs further investigation.

The decrease in root hydraulic conductance in dry soil may also be attributed to root shrinkage or reduced apoplastic water uptake. While mucilage and root hairs aid in maintaining cohesion between the root and surrounding soil, root hair shrinkage can occur at soil water potentials below -10 kPa, followed by cortex shrinkage, leading to air gaps at the root-soil interface (Ahmed et al. 2018a; Duddek et al. 2022; Jiang et al. 2022). Our results indicated potential shrinkage in crown roots – in wet soil, the cortex of crown roots was 43 – 65% thicker than primary roots and seminal roots; however, in dry soil, the cortex of 1^st^ whorl crown roots decreased to a level similar to that of primary and seminal roots (Fig.S7) as reported earlier (Carminati et al. 2013). Shrinkage at the V6 stage was not observed, probably because maize with higher-order crown roots manages to absorb sufficient water for growth. Moreover, water uptake may partially shift from the apoplastic pathway to the cell-to-cell pathway in dry soil as indicated by the decreased transpiration due to partial stomatal closure (Fig. S8) (Suslov et al. 2024). The thicker cortex layer of crown roots might retard the radial water movement because the cell-to-cell pathway is 17 times slower than the apoplastic pathway (Steudle 2001). Therefore, future research should investigate root shrinkage and water transport pathways across different root types under water deficit conditions, as these factors significantly impact water and nutrient uptake.

Under wet soil, crown roots exert greater hydraulic conductance than primary and seminal roots (Fig. 2). On average, crown roots of V3 maize had hydraulic conductivity of 6 ×10^-11^ m^3^ Mpa^-1^ s^-1^ and primary/seminal roots had on average 1.8 ×10^-11^ m^3^ Mpa^-1^ s^-1^, both falling within the range of 0.6 ×10^-11^ – 6 ×10^-11^ m^3^ Mpa^-1^ s^-1^ (Knipfer and Fricke, 2011) using the root exudation method. Greater hydraulic conductance in crown roots is attributed to their larger, numerous xylem vessels than in primary and seminal roots because plants translocate water and NO_3_^-^ in the xylem. For instance, a single crown root produced seven metaxylem vessels, whereas seminal roots only produced four metaxylem vessels (Table 1). The number of xylem vessels in the present study is smaller than the B73 maize inbred line, which contains 9–12 xylem vessels in the crown roots and 4–6 xylem vessels in the primary and seminal roots (Tai et al. 2016). However, the xylem diameter in crown roots of the B73 maize inbred line was 27–68% larger than in primary and seminal roots (Tai et al. 2016), aligning with our results. Thus, the larger xylem vessel in crown roots produces a higher tension that drives the NO_3_^-^ mass flow to the root surface. A greater difference in measured root hydraulic conductance than the calculated axial hydraulic conductance among the root types indicates that crown roots also have radial hydraulic conductance than embryonic roots, possibly due to the activity of aquaporins (Chaumont and Tyerman 2014). Furthermore, maize crown root tips exhibited higher NO_3_^-^ uptake kinetics compared to seminal root tips, with a higher (> 20%) maximum NO_3_^-^ influx rate (York et al. 2016). Therefore, the crown roots dominate NO_3_^-^ uptake because of their high hydraulic conductance and the activity of their abundant NO_3_^-^ transporters on the root surface. Further study is required to confirm this possibility.

Our results suggested no significant difference in soil nitrification among root types, contrary to our initial hypothesis. Nitrification (a biological process controlled by ammonium oxidizers and nitrifiers) is influenced by soil physio-chemical properties such as soil moisture and pH. In the present study, the nitrification rates ranged from 18 – 33 mg NO_3_^-^ kg^-1^ d^-1^ in the wet soil and 12 – 20 mg NO_3_^-^ kg^-1^ d^-1^ in the dry soil (Fig. 4), falling within the reported range of < 1 – 50 mg NO_3_^-^ kg^-1^ d^-1^ (Lteif et al. 2010; Zhang et al. 2024). The elevated nitrification rate around crown roots in the wet soil than the dry soil was consistent with the optimal soil moisture range for nitrification (-5 – -20 kPa, equivalent to 50 % – 80% of field capacity) as reported earlier (Whalen and Sampedro, 2010). Notably, embryonic roots exhibited higher pH than crown roots (Fig. S9) likely because crown roots exude more organic acids (Tiziani et al. 2022). Nevertheless, soil pH remained within the range of 5.8 – 7.0, which favors the nitrification of autotrophic nitrifiers and results in uniform nitrification among the root types (Whalen and Sampedro 2010). Further investigation into how soil N transformation varies across diverse root types will contribute to a precise understanding of soil-plant interactions.

In summary, N uptake in maize depends on the development of crown roots, which have greater hydraulic conductance than embryonic roots. Under dry soil conditions, N uptake and crown root hydraulic conductance decreased at the V3 and V6 stages. The embryonic roots can contribute to maize N uptake in early vegetative growth (up to the V3 stage), but their water and NO_3_^-^-absorbing functions diminish by the V6 stage as the larger, solute-extracting crown root system develops. The present results suggest that soil water management is important to ensure optimal N uptake, especially at the V6 stage and thereafter because the embryonic root functions are negligible.

## Author contributions

YJ and JW conceived the research. YJ performed the experiments, analyzed, interpreted the data, and wrote the manuscript. JW participated in data interpretation and revised the manuscript. JW supervised the work. All authors approved the manuscript.

## Acknowledgments

We appreciate technical support from Youssef Chebli, Mohammed Antar, Sarah-Ann Persechino, Emily Ngo Schick, Khosro Mousavi, and Aurélie Stil. We are grateful to Dr. Chih-Yu Hung for his contribution to the early experimental design stage. We thank Dr. Geoffrey Sunahara for his kind review of an earlier version of this paper.

## Data availability

The data generated and analyzed during the current study are provided, in part, as supplementary material and are available from the corresponding author upon reasonable request.

## Supplementary information

The online version contains supplementary material available at (weblink to be generated by publisher).

